# Automatic variable selection in ecological niche modeling: A case study using Cassin’s Sparrow (*Peucaea cassinii*)

**DOI:** 10.1101/2021.09.03.458913

**Authors:** John L. Schnase, Mark L. Carroll

**Affiliations:** Office of Computational and Information Sciences and Technology, NASA Goddard Space Flight Center, Greenbelt, Maryland, 20708, USA

## Abstract

MERRA/Max provides a feature selection approach to dimensionality reduction that enables direct use of global climate model outputs in ecological niche modeling. The system accomplishes this reduction through a Monte Carlo optimization in which many independent MaxEnt runs, operating on a species occurrence file and a small set of randomly selected variables in a large collection of variables, converge on an estimate of the top contributing predictors in the larger collection. These top predictors can be viewed as potential candidates in the variable selection step of the ecological niche modeling process. MERRA/Max’s Monte Carlo algorithm operates on files stored in the underlying filesystem, making it scalable to large data sets. Its software components can run as parallel processes in a high-performance cloud computing environment to yield near real-time performance. In tests using Cassin’s Sparrow (*Peucaea cassinii*) as the target species, MERRA/Max selected a set of predictors from Worldclim’s Bioclim collection of 19 environmental variables that have been shown to be important determinants of the species’ bioclimatic niche. It also selected biologically and ecologically meaningful predictors from a more diverse set of 86 environmental variables derived from NASA’s Modern-Era Retrospective Analysis for Research and Applications Version 2 (MERRA-2) reanalysis, an output product of the Goddard Earth Observing System Version 5 (GEOS-5) modeling system. We believe these results point to a technological approach that could expand the use global climate model outputs in ecological niche modeling, streamline the modeling process, and, eventually, enable automated bioclimatic modeling as a practical, readily accessible, low-cost, commercial cloud service.

## Introduction

Ecological niche modeling (ENM) consists of a set of techniques and tools that use species occurrence records and environmental data to predict the relative suitability of habitats [1]. It is used across a wide range of disciplines, including fields as diverse as biogeography and phylogeny [2], conservation biology and epidemiology [3,4], invasion biology [5], and archaeology [6]. In recent years, ecological niche models have become particularly important in understanding the influence of climate change on the geographic distribution of species [7]. This, in turn, has led to greater use of global climate model (GCM) outputs as environmental predictors [8]. GCMs provide global representations of the climate system, projections for hundreds of variables, and combine observations from an array of satellite, airborne, and *in-situ* sensors [9]. The largest and most sophisticated of these, however, produce complex, petabyte-scale data sets, which complicates variable selection and limits their direct use in ecological modeling [10–12].

In previous work, we demonstrated the potential of a MaxEnt-based Monte Carlo method that addresses this issue by screening large data collections for viable predictors [13]. Based on a machine learning approach to maximum entropy modeling, MaxEnt is one of the most popular software packages in use today by the ENM community [14–16]. Among its many advantages, MaxEnt ranks the contribution of predictor variables in the formation of its models. Our Monte Carlo method exploits this feature in an ensemble strategy whereby many independent MaxEnt runs, each drawing on a small, random subset of variables stored in the filesystem, converge on a global estimate of the top contributing subset of variables in the larger collection. These top-contributing predictors can then be studied in more detailed ways, augmented with other variables, and further refined prior to final model construction. We believe a screening step, such as this, could help the ENM process, particularly when working with large, multidimensional data sets where selection through ecological reasoning or other means is not apparent.

In our earlier, proof-of-concept work, we implemented the Monte Carlo selection algorithm as a single-threaded program running on a MacBook Pro laptop computer [13]. In the current study, we have implemented a parallel version of the Monte Carlo method in a high-performance cloud computing environment. Our goal this time has been to characterize the run-time performance and scaling properties of a parallel implementation of the Monte Carlo algorithm and demonstrate its variable selection behavior with two example use cases. We call the prototype system MERRA/Max to reflect its reliance on MaxEnt and our interest in using the technology to screen for bioclimatic predictors in NASA’s Modern-Era Retrospective Analysis for Research and Applications Version 2 (MERRA-2) dataset, what we view as an underutilized and potentially important GCM resource for the ecological modeling community.

MERRA/Max’s parallel implementation scales linearly with respect to the variable sampling rate. Because each MaxEnt sampling run is independent of all other runs in an ensemble, MERRA/Max can screen an entire data collection in the time it takes to complete a single MaxEnt run when sufficient processors are available for complete parallelism. In a use case scenario involving Cassin’s Sparrow (*Peucaea cassinii*) as the target species and Worldclim’s classic set of 19 Bioclim variables as the collection to be scanned, MERRA/Max identified a set of predictors known from the published literature to be important determinants of the species’ bioclimatic niche. In a second scenario involving a novel collection of 86 MERRA-2 variables, MERRA/Max likewise identified as top predictors a set of variables with biological and ecological relevance to Cassin’s Sparrow. Taken together, these results suggest that MERRA/Max and its Monte Carlo approach to variable selection may provide an effective way of screening large collections of environmental variables for potential predictors, thereby contributing to the variable selection step of the ENM process.

This project builds on a twenty-year history of technology research and development at NASA focusing on applications of high-performance computing to ecological modeling [17–21] and the big data challenges of Earth science [22–32]. It complements this body of work by looking at ways that machine learning and high-performance cloud computing can extend existing capabilities and open new opportunities for research. In a fully realized, operational implementation of the technologies described here, we see MERRA/Max as one element of a bioclimatic modeling service enabled by a suite of high-performance data subsetting and data analytic tools of the sort becoming increasingly available to the research community through commercial cloud services [33–38].

## Materials and Methods

### System architecture and implementation

We implemented MERRA/Max in a 100-core testbed within the NASA Center for Climate Simulation’s (NCCS’s) Advanced Data Analytics Platform (ADAPT). ADAPT is a managed virtual machine (VM) environment most closely resembling a platform-as-a-service (PaaS) cloud [39]. It features over 300 physical hypervisors that host one or more VMs, each having access to multiple shared, centralized data repositories. The hypervisor hardware consists of 2.2 GHz 24-core Intel Xeon Broadwell E5-2650 v4 processors with 256 GB of memory. The MERRA/Max testbed consists of a dedicated set of ten 10-core Debian Linux 9 Stretch VMs. We used shell scripts, R Version 4.0.1 [40], ENMeval Version 0.3.1 [41], and MaxEnt Version 3.4.1 [42] to develop MERRA/Max’s software components, which collectively realize the Monte Carlo algorithm through the interactions shown in Fig 1.

**Fig 1.**
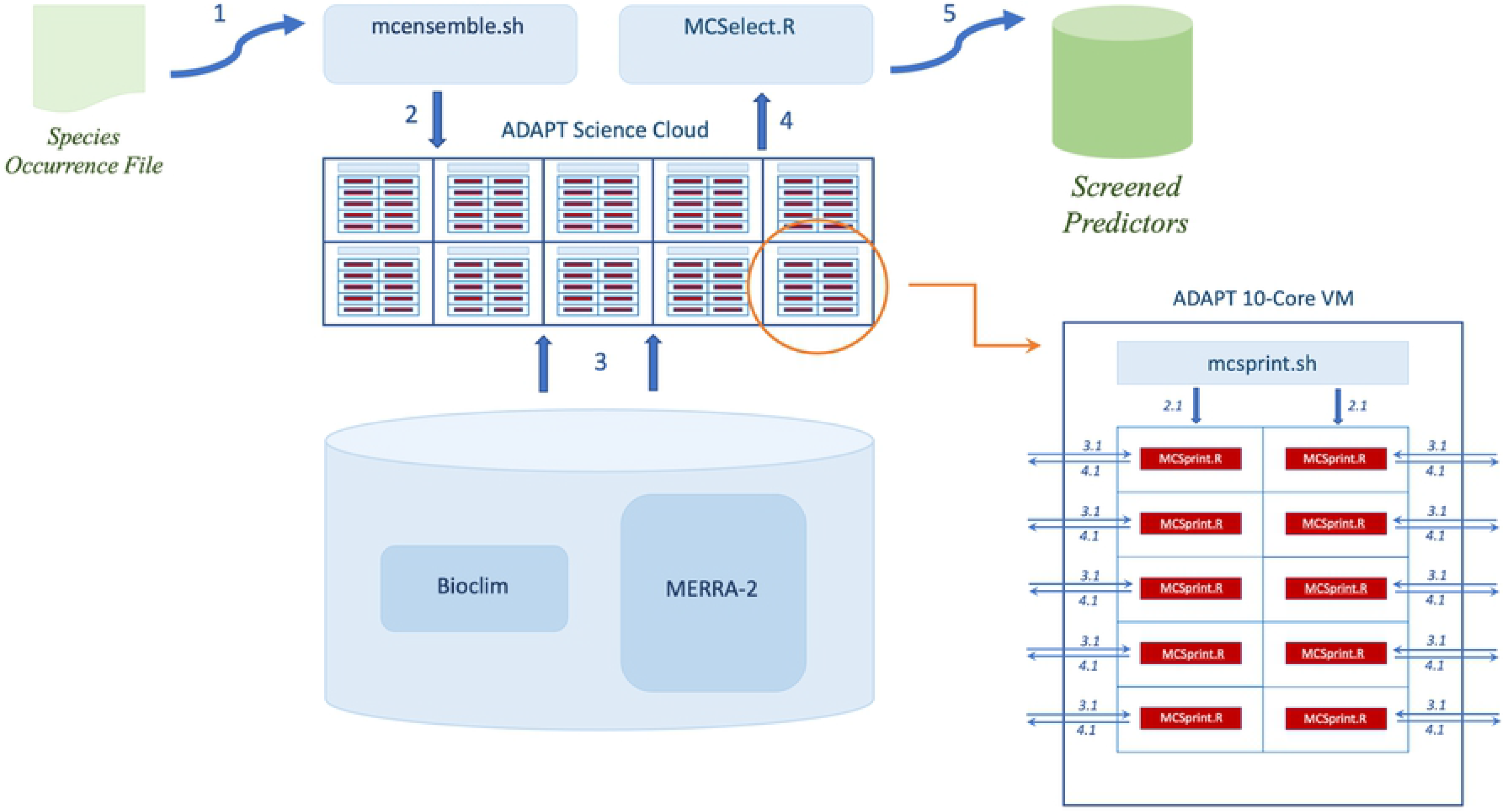
MERRA/Max architecture. Conceptual diagram showing the major hardware and software components of the MERRA/Max prototype. The study’s testbed consisted of 10 virtual machines (VMs) within NASA’s ADAPT science cloud, with each VM contributing 10 processing cores to the testbed. Numbered arrows indicate the system’s processing workflow.

Conceptually, MERRA/Max sits atop a collection of variables stored in the underlying filesystem; when provided a species occurrence file, the system screens the collection to find the most important MaxEnt predictors for the input provided (Fig 1, Steps 1–5). The Monte Carlo screening process is initiated by the *mcensemble.sh* script, which launches an *mcsprint.sh* script on each of the 10 MERRA/Max VMs (Fig 1, Step 2). The *mcsprint.sh* script, in turn, creates parallel sprint runs by launching an R run-time environment and an *MCSprint.R* program on each of the 10 VM’s 10 processor cores (Fig 1, Steps 2.1). The *MCSprint.R* programs perform repeated MaxEnt runs on random pairs of variables read from the shared filesystem until a desired level of sampling is achieved (Fig 1, Steps 3.1). *MCSprint.R* maintains a tally table that tracks the number of times each variable is used along with its accumulating permutation importance, then writes the table to a shared directory (Fig 1, Steps 4.1). We used the operating system’s *MCSprint.R* process identifiers to create unique *pid.tt* file names for the output tally tables. When all the sprints have completed their work, *MCSelect.R* concatenates the *pid.tt* files into a global tally table (Step 4), computes the average permutation importance for each variable, then sorts them to reveal the top contributing variables identified by the ensemble’s runs (Fig 1, Step 5). While MERRA/Max relies on MaxEnt to perform selection, it is important to note that the resulting set of selected variables can be used in any ENM application or species distribution modeling approach.

### Run-time performance and scaling properties

MERRA/Max’s run-time performance and scaling properties are affected by several factors. The total amount of time needed for MERRA/Max to complete a screening run (T) is primarily determined by the number of variables in the collection being screened (N), the average number of random samples taken of each variable in the collection (S), the number of random variables used in each independent MaxEnt sampling run (V), and the number of processor cores available in the compute environment (C). To understand the interplay of these factors, we gathered timing metrics on a series of ensembles with varying values of N, S, and C.

In a first ensemble with N = 2 and C = 10, we used 10 *parallel* sprints (one sprint per core) in which each sprint performed five *sequential* two-variable sampling runs to achieve an average sample size for each collection variable of S = 50. We then completed the S = 50 series with ensembles in which N and C were proportionally increased to N = 18 and C = 90. This process was repeated using 10- and 15-run sprints respectively to create a timing series for S = 100 and S = 150. These measurements allowed us to quantify MERRA/Max’s run-time performance, estimated optimal performance, and its scaling properties within the constraints of a 100-core testbed.

We did not quantify non-algorithmic influences on run time, such as predictor resolution, occurrence file size, competing system processes, filesystem performance, processor failures, process failures (abends), or MaxEnt parameter settings, as these tend to be intrinsic properties of the science question being studied, the compute environment, or the MaxEnt software itself. These non-algorithmic factors either have an idiosyncratic impact on overall run time that is constant for any particular application of MERRA/Max, or they are beyond user control.

For development testing, we used Cassin’s Sparrow (*Peucaea cassinii* Woodhouse, 1852) as the target species [43] and Worldclim Version 2.1’s 19 Bioclim variables at a resolution of 5.0 arc-minutes (Table 1) as environmental predictors [44,45]. We obtained Cassin’s Sparrow observational records for the year 2016 from the Global Biodiversity Information Facility (GBIF) [46]. Because of their secretive nature, Cassin’s Sparrows are generally detected in the field by the presence of singing males that define and defend breeding territories that range in size from 0.6 to 12.9 acres [43,47–49]. After removing replicates, we thinned the records to non-overlapping observations within a 16 km buffer around each point to avoid double counting the same individuals. This resulted in a total of 609 observations, which were used for testing throughout the study. The predictor layers were clipped to the coverage area of our observational data, reprojected, and formatted for use by MaxEnt using rgdal Version 1.5-18 [50] following the guidelines of Hijmans et al. [51].

**Table 1.** Bioclim variables.

|  |  |
| --- | --- |
| bio01 | Annual mean temperature |
| bio02 | Mean diurnal range (mean of monthly (max temp - min temp)) |
| bio03 | Isothermality (bio2/bio7) ($\times 100$) |
| bio04 | Temperature seasonality (standard deviation $\times 100$) |
| bio05 | Maximum temperature of warmest month |
| bio06 | Minimum temperature of coldest month |
| bio07 | Temperature annual range (bio5-bio6) |
| bio08 | Mean temperature of wettest quarter |
| bio09 | Mean temperature of driest quarter |
| bio10 | Mean temperature of warmest quarter |
| bio11 | Mean temperature of coldest quarter |
| bio12 | Annual precipitation |
| bio13 | Precipitation of wettest month |
| bio14 | Precipitation of driest month |
| bio15 | Precipitation seasonality (coefficient of variation) |
| bio16 | Precipitation of wettest quarter |
| bio17 | Precipitation of driest quarter |
| bio18 | Precipitation of warmest quarter |
| bio19 | Precipitation of coldest quarter |

We adopted a standard MERRA/Max screening configuration that we used as the default in all our timing trials and use cases. This included a MaxEnt feature class (FC) setting of LQHP (linear, quadratic, hinge, and product), a regularization multiplier (RM) setting of 1.0, 10 replicate cross-validation, and ten thousand background points from across the study area [13]. We used V = 2 random variables in all the independent MaxEnt sampling runs. Additional detail about MERRA/Max’s default screening parameters and the rationale for their choice are provided as supporting information in S1 Appendix.

### Use case scenarios and selection behavior

To demonstrate MERRA/Max’s selection behavior and show how the system might be used in actual practice, we developed two use case scenarios in which we modeled the bioclimatic niche of Cassin’s Sparrow, a species known to be sensitive to many of the variables used in the study [43,47,49,52]. Each use case involved three steps. The first was a *Variable Screening* step, in which MERRA/Max selected the top six contributing predictors from a collection of variables using an average sampling rate of S = 50. We used the averaged results from three screenings to settle on the top contributors. This was followed by a *Predictor Refinement* step, where we used variance inflation factor (VIF) analysis to reduce collinearities in the selected predictors [53]. VIF shows the degree to which standard errors are inflated due to the levels of multicollinearities. Using ENMtools Version 1.4.4 [54], we first calculated Pearson correlation coefficient (r), coefficient of determination (r^2^), and VIF [1÷ (1–r^2^)] values for the selected predictors, then eliminated the least contributing variable in any pair of variables having r > 0.8, r^2^ > 0.8, and VIF > 10.0 [53]. In a final *Model Calibration / Final Model Run* step, we used the ENMeval R package [41,55] to identify optimal settings for the remaining, non-collinear predictors by performing a series of MaxEnt runs across all possible combinations of five feature classes (L, LQ, H, LQH, and LQHP) and regularization multiplier values ranging from 0.5 to 4.0 in increments of 0.5. The combination of settings resulting in the lowest value for Akaike’s information criterion corrected for small sample size (AICc) [56] was taken to be an optimal tuning configuration.

We used the same 2016 Cassin’s Sparrow occurrence data in each scenario that we used for development testing. However, the two use cases operated on different sets of environmental predictors. In the first, we again used WorldClim’s 19 Bioclim variables. In the second use case, to gain experience with an even larger collection and demonstrate the system’s application to a novel set of Intergovernmental Panel on Climate Change (IPCC)-class GCM outputs [10,57], we used variables obtained directly from the Modern-Era Retrospective Analysis for Research and Applications Version 2 (MERRA-2) reanalysis. In contrast to Worldclim’s Bioclim predictors, which are derived from spatially interpolated weather station temperature and precipitation data, the MERRA-2 reanalysis is produced by NASA’s Goddard Earth Observing System Version 5 (GEOS-5) [58–60]. The system integrates observational data with numerical models to produce a global temporally and spatially consistent synthesis of over 600 climate-related variables. MERRA-2’s spatial resolution is 1/2 latitude × 5/8 longitude (i.e., 55.5 × 69.4 km at the equator) × 72 vertical levels extending through the stratosphere. Its temporal resolution is hourly and extends from 1979 to the present, nearly the entire satellite era. The complete MERRA-2 collection is about one petabyte in size.

For the current study, we created a test collection of 86 MERRA-2 variables of potential ENM interest. These were drawn from four MERRA-2 collections and included modeled, two-dimensional values for atmospheric attributes and heat, wind, radiation, and land surface attributes (Table 2). The test collection contains weekly and monthly maximum, minimum, and average values (or sums as appropriate) for each variable for the 40 years spanning 1980 to 2020. Importantly, the collection contains modeled values for the temperature and precipitation variables that form the basis for Bioclim’s 19 predictors, which highlight climate conditions generally understood to relate to a species’ physiology, *plus* an extended array of environmental attributes of potentially more direct biological significance, such as soil moisture and evaporation, wind direction and speed, and various solar radiation fluxes (Table 2) [44,61–65]. For our use case, we used xarray [66] to create annual averages for the 86 variables for the year 2016, the year corresponding to the observation year of our Cassin’s Sparrow occurrence data. We used modeled values at 850 hPa, where appropriate, to reflect surface conditions. The hectopascal (hPa) atmospheric pressure unit is an expression of altitude. Generally, 850 hPa lies immediately above the atmospheric boundary layer (about 1.5 km), where daily surface variations in temperature, humidity, wind speed, etc. have little if any effect on measured or modeled values [9]. These layers were then prepared for use with MaxEnt as described above.

**Table 2.**
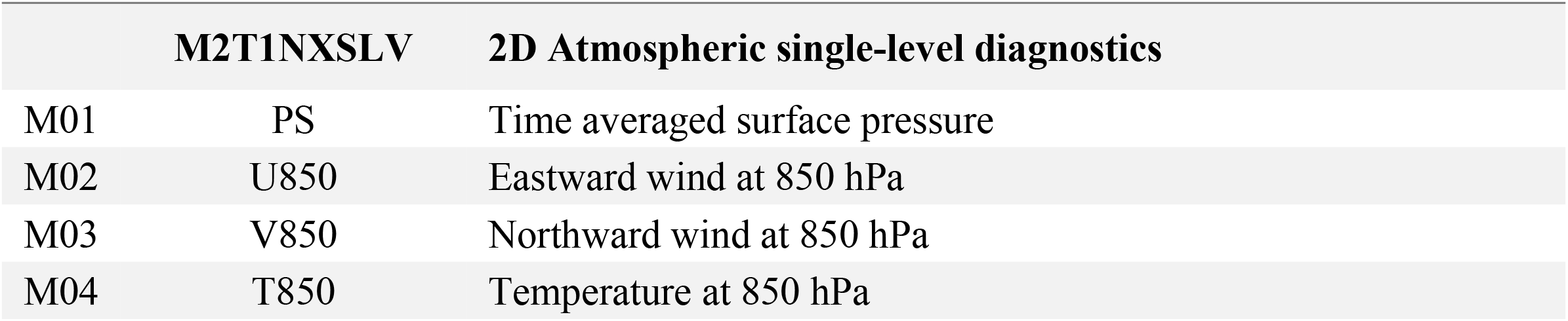
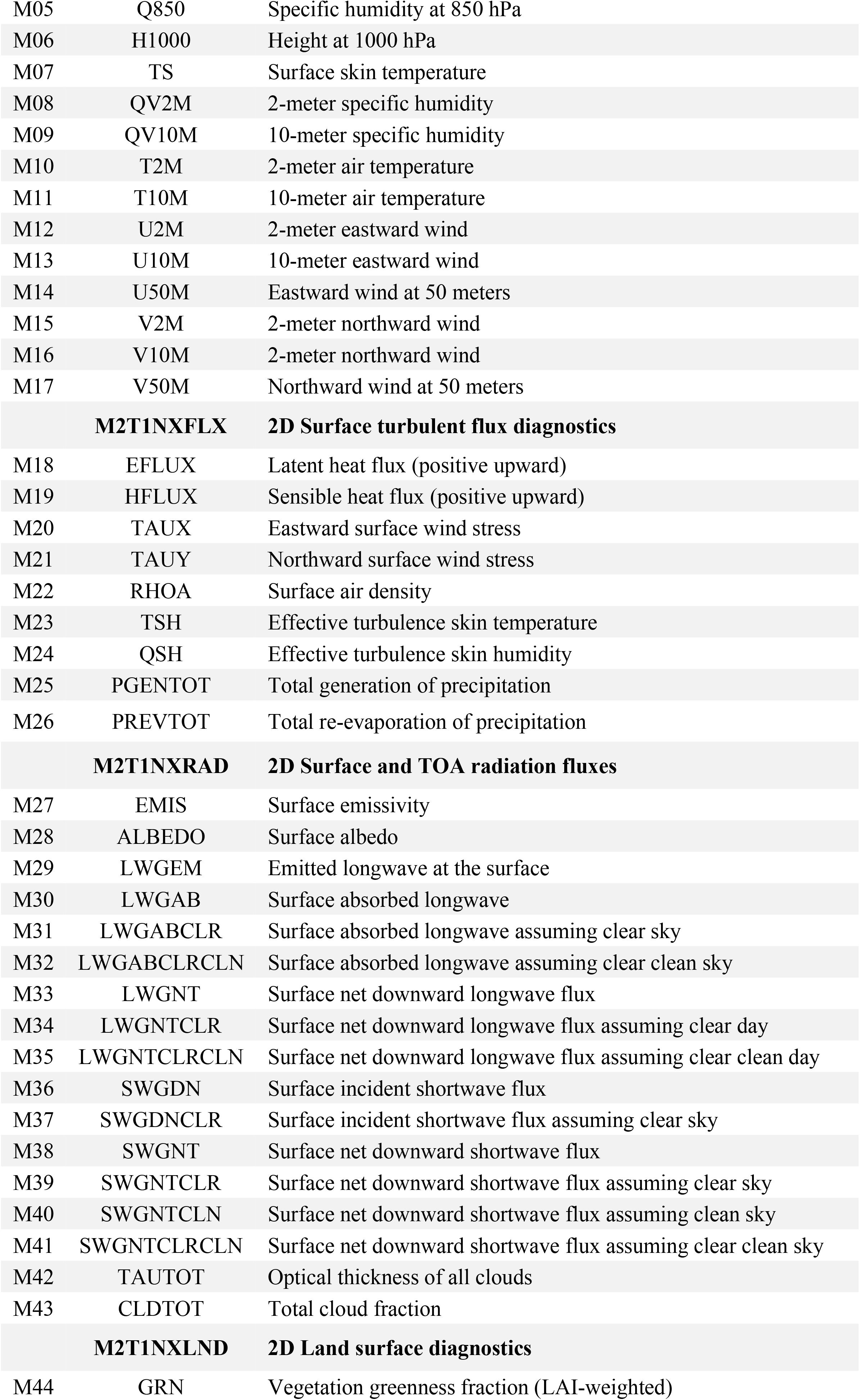
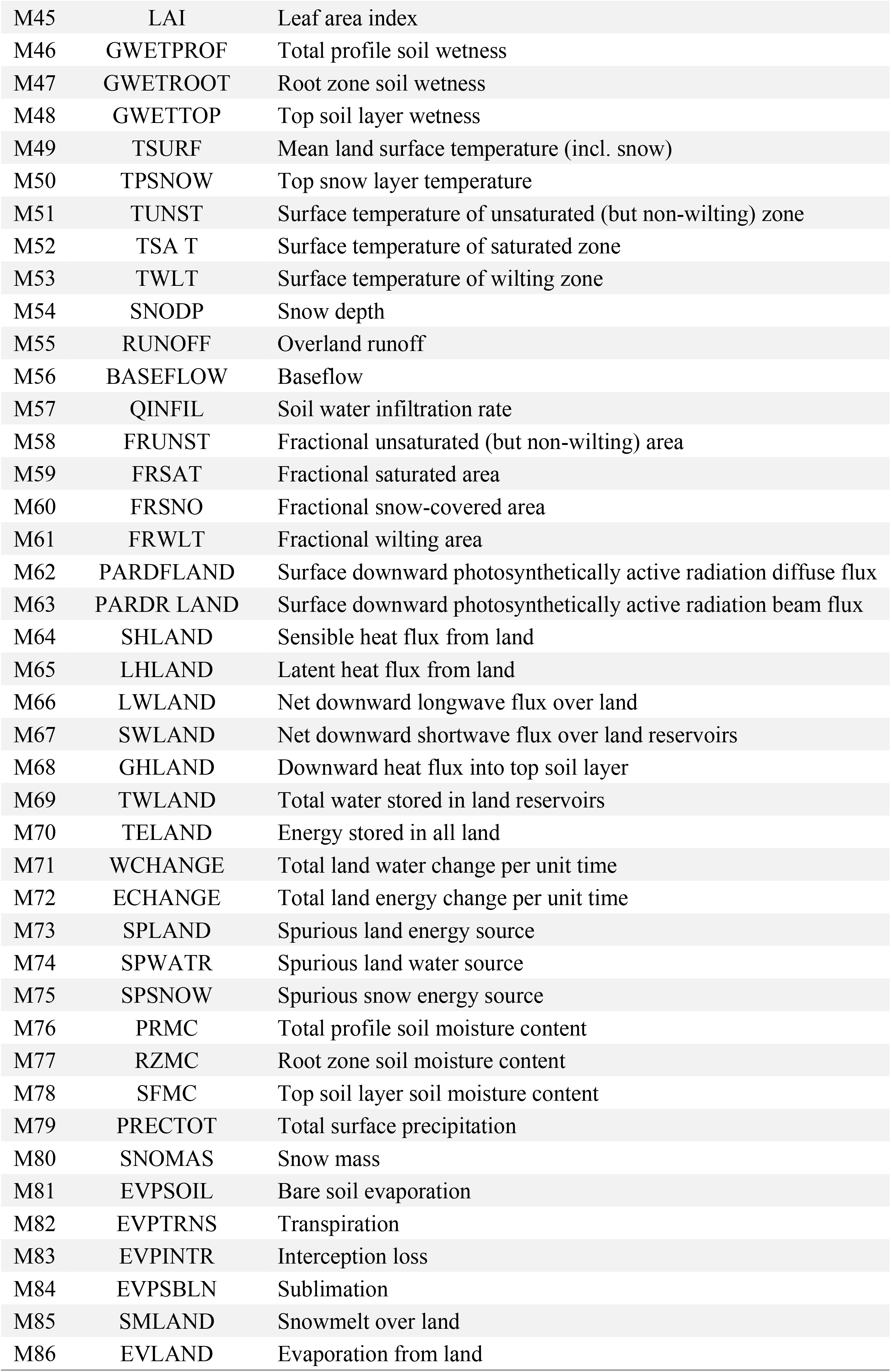
MERRA-2 variables.

To evaluate MERRA/Max’s selection behavior, we created initial MaxEnt models using the top six predictors selected by the three screenings in the *Variable Screening* step. Then, using the overall top six variables found in the *Variable Screening* step, we created a final MaxEnt model in the *Model Calibration / Final Model Run* step that reflected any improvements gained in the *Predictor Refinement* step or by *Model Calibration*. The potential distribution maps produced by the final models were judged for reasonableness based on first-hand knowledge of the species, its habitat preferences, what is known about Cassin’s Sparrow’s range from the published literature [43,47–49,52], and observational records from Cornell Lab’s eBird citizen-scientist database [67].

We further compared these final model predictions to results obtained by replicating, in part, the work of Salas et al. [68], in which traditional MaxEnt variable-selection techniques were used to model the bioclimatic niche of Cassin’s Sparrow. Here, we used our 2016 Cassin’s Sparrow occurrence data in combination with the seven Worldclim Bioclim variables used by the Salas team: bio03, bio06, bio08, bio09, bio12, bio14, and bio18. The Salas team chose these predictors by first removing one of each pair of highly correlated variables to avoid collinearity among the variables. The team then chose between highly correlated variables by selecting those that were identified in one or more species-specific studies as having an effect on the species’ range or population dynamics. In cases where the literature search could not differentiate between two highly correlated variables, the team used a qualitative assessment of the distribution of values of the variable at all presence points and the relationship between the variable and species presence or pseudo-absence [68]. We used ENMeval, as described above, to identify optimal tuning parameters for the Salas-derived model.

To gain a quantitative perspective on performance, we used AICc [56] as a measure of a model’s relative explanatory power (lower values indicating less information loss) and area under the receiver operating characteristic curve (AUC) [69], percent correctly classified (PCC) [70], and the True Skill Statistic (TSS) [71,72] as measures of model accuracy (higher values in all cases indicating greater accuracy). Similarities between our first use case’s final Bioclim model and the Salas-derived Bioclim model were examined using Warren’s *I* statistic [73], Schoener’s *D* statistic [74], and Pearson’s *r* statistic [75]. All input data used in this study, along with a set of example scripts are provided as supporting information in S2 File.

## Results

### Run-time performance and scaling properties

The first ensemble of the 50-sample timing series (S = 50) required a total run time of T = 7.9 minutes to screen a two-variable collection (N = 2) using 10 processor cores (C = 10) (Fig 2A). At one sprint per core, and with each MaxEnt sampling run operating on two (V = 2) randomly selected variables at a time, this first ensemble needed 50 MaxEnt runs to do its work. The shortest achievable screening time (T*min*) is possible only when the number of cores needed for perfect parallelism (C*max*) are actually available, in this case, 50:

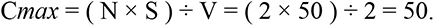

**Fig 2.**
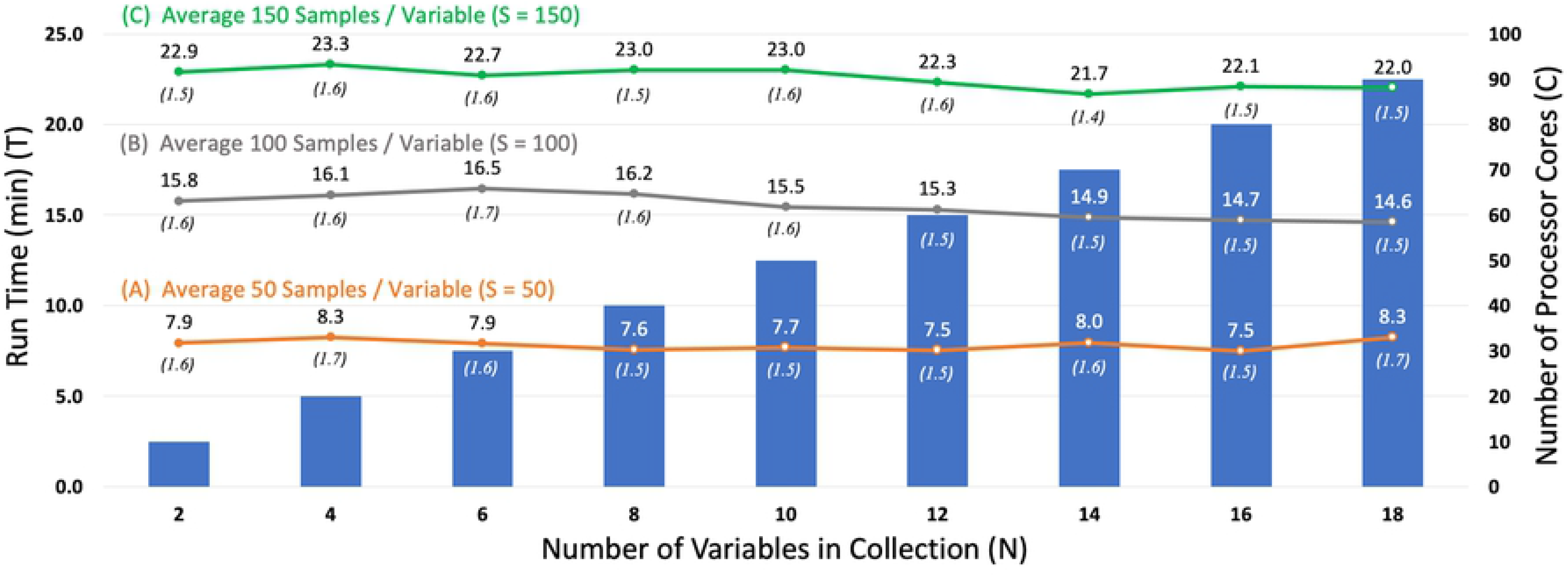
MERRA/Max run-time performance and scaling properties. Figure shows the relationship between the amount of time it takes MERRA/Max to complete a screening run (T) (shown by the colored lines and left Y axis), the number of variables in the collection being scanned (N), the average number of random samples taken of each variable in the collection during the screening process (S), and the number of processor cores available in the compute environment (C) (shown by the colored vertical bars and right Y axis. MERRA/Max’s parallel implementation scales linearly with respect to S, and, for any given collection of size N and sample size S, the estimated minimum possible run time (T*min*) (shown in parentheses) can be achieved when enough cores are available for a completely parallel screening of the collection.

Because only 10 cores were available, each of the parallel sprints had to perform five sequential MaxEnt sampling runs to achieve the S = 50 sampling goal, a repeat factor (R) of 5:

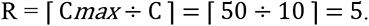

By accounting for this performance cost, we estimate that MERRA/Max’s minimum possible run time, in a completely parallel screening of this first data set, would have been about 1.6 minutes:

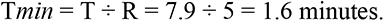

In each subsequent ensemble of the S = 50 series, we added two variables to the scanned collection and 10 cores to the pool of available processors. With this proportional scaling of variables and processors, average run times remained constant across the series at T = 7.9 ± 0.3 minutes (T*min* = 1.6 ± 0.1 minutes) (Fig 2A). In the S = 100 timing series, R = 10 sequential MaxEnt runs were used in each sprint to achieve the desired sampling level (Fig 2B), and in the S = 150 series, R = 15 runs were used (Fig 2C). In both cases, run times scaled linearly with sample size and remained relatively constant across the series, with T = 14.8 ± 2.5 minutes (T*min* = 1.6 ± 0.1 minutes) for the S = 100 series and T = 22.6 ± 0.5 minutes (T*min* = 1.5. ± 0.1 minutes) for the S = 150 series.

### Use case scenarios and selection behavior

The Bioclim collection consists of N = 19 variables. To achieve an average per-variable sampling goal of S = 50 with C = 100 cores, each sprint in the Bioclim use case (Fig 3A) performed R = 5 sequential MaxEnt runs in the *Variable Screening* step, resulting in ensembles comprising a total of 500 runs. In an average of three such ensembles, MERRA/Max took T = 6.4 ± 0.5 minutes (T*min* = 1.3 ± 0.1 minutes) to identify bio18, bio03, bio05, bio08, bio13, and bio16 as the top six contributing variables of the collection. In the subsequent *Variable Refinement* step, predictor pairs bio13-bio16 and bio16-bio18 were shown to be correlated, which led us to discard bio16 from the selection set. In the *Model Calibration / Final Model Run* step, the remaining five non-correlated variables were used to create a final model in which the top four contributing variables (bio18, bio03, bio05, and bio13) accounted for approximately 98% of overall permutation importance, and the performance metrics were AICc 12,232, AUC 0.83, PCC 0.75, and TSS 0.49.

**Fig 3.**
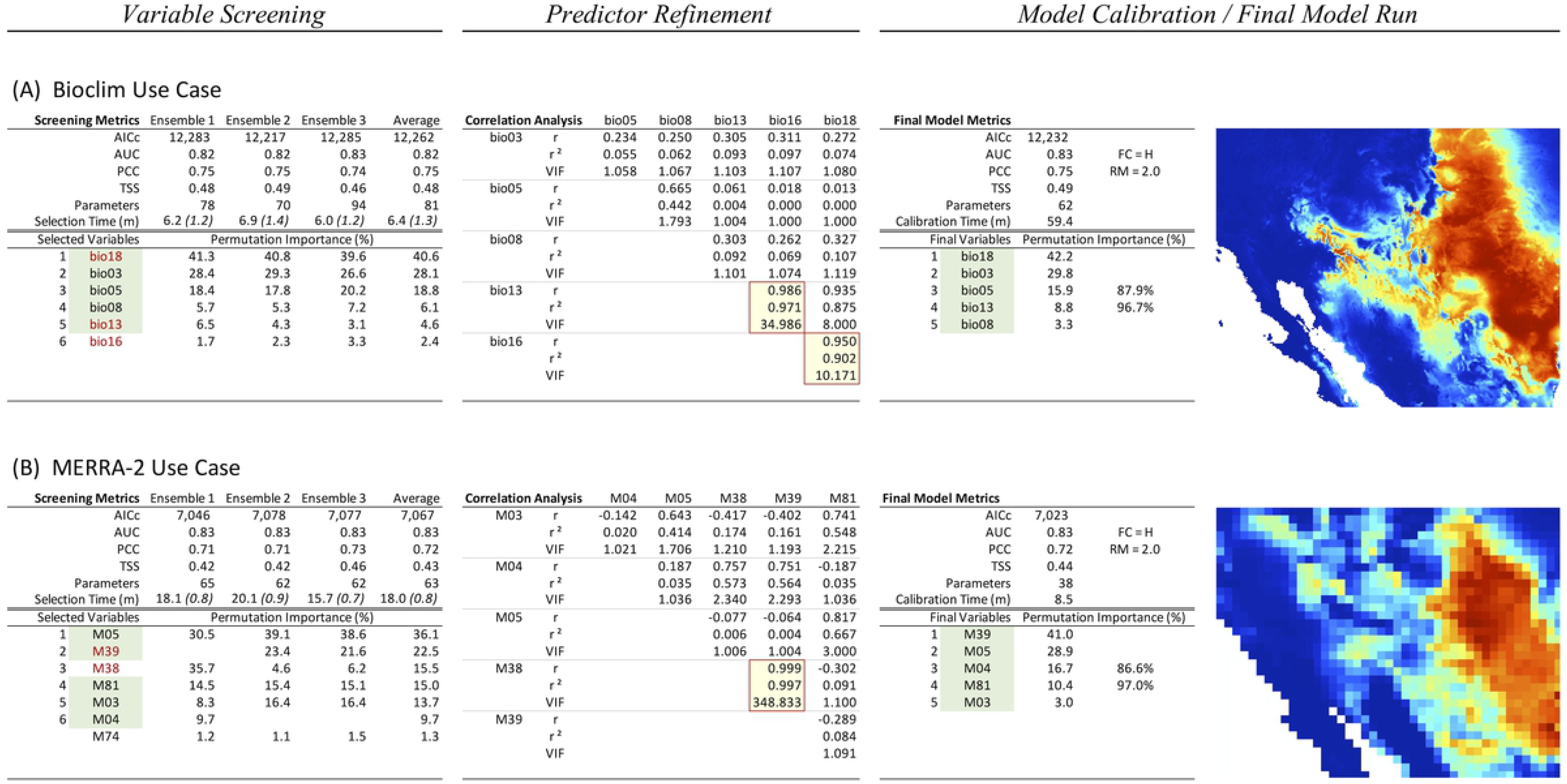
MERRA/Max use case scenarios. Figure shows the results of two use cases involving Cassin’s Sparrow observational data and predictor data sets of contrasting size and complexity: the Bioclim collection with N = 19 variables (A) and a MERRA-2 reanalysis test collection comprising N = 86 variables (B). A *Variable Screening* step was used in each scenario to select the top six contributing variables in the underlying collection. Correlated variables (indicated with red text and yellow highlight) were identified in a *Predictor Refinement* step and thinned to reduce collinearities. In a third step, *Model Calibration* and a *Final Model Run* were performed with the remaining non-correlated variables (green highlight). AICc is Akaike’s information criterion corrected for small sample size, AUC is area under the receiver operating characteristic curve, PCC is percent correctly classified, TSS is True Skill Statistic, Parameters is MaxEnt’s measure of model complexity, r is Pearson’s correlation coefficient, r^2^ is the coefficient of determination, and VIF is variable inflation factor. The estimated minimum run time (T*min*) for a completely parallel screening is shown in parentheses. Maps created by the authors show MaxEnt logistic output, which can be interpreted as an estimated probability of presence between 0 and 1 with warmer colors indicating better predicted conditions for the species.

In the MERRA-2 use case (Fig 3B), MERRA/Max screened a collection of N = 86 variables of coarser resolution (approximately 50 km for MERRA-2 vs. 8 km for Bioclim). To achieve the S = 50 sampling goal, each sprint performed R = 22 MaxEnt sampling runs, which resulted in 2200-run ensembles. The average run time across three such ensembles in the *Variable Screening* step increased to T = 18.0 ± 2.2 minutes; however, because of the coarser predictor resolution, times for the MaxEnt sampling runs decreased, which resulted in an estimated theoretical lower bound of only T*min* = 0.8 ± 0.1 minutes. The six top contributing variables identified in the *Variable Screening* step included M05, M39, M38, M81, M03, and M04. In the *Predictor Refinement* step, the M38-M39 pair showed strong correlation, which led us to discard M38. In the *Model Calibration / Final Model Run* step, the remaining five non-correlated variables were used to create a final model in which the top four contributing variables (M39, M05, M04, and M81) accounted for approximately 97% of over overall permutation importance, and the performance metrics were AICc 7,023, AUC 0.83, PCC 0.72, and TSS 0.44.

In the Salas-derived Cassin’s Sparrow model (Fig 4A), where a traditional approach to variable selection was used to identify the seven predictors used in MaxEnt, the top four contributing variables (bio18, bio06, bio14, and bio09) accounted for approximately 83% of overall permutation importance, and the model’s performance metrics were AICc 12,169, AUC 0.83, PCC 0.76, and TSS 0.50.

**Fig 4.**
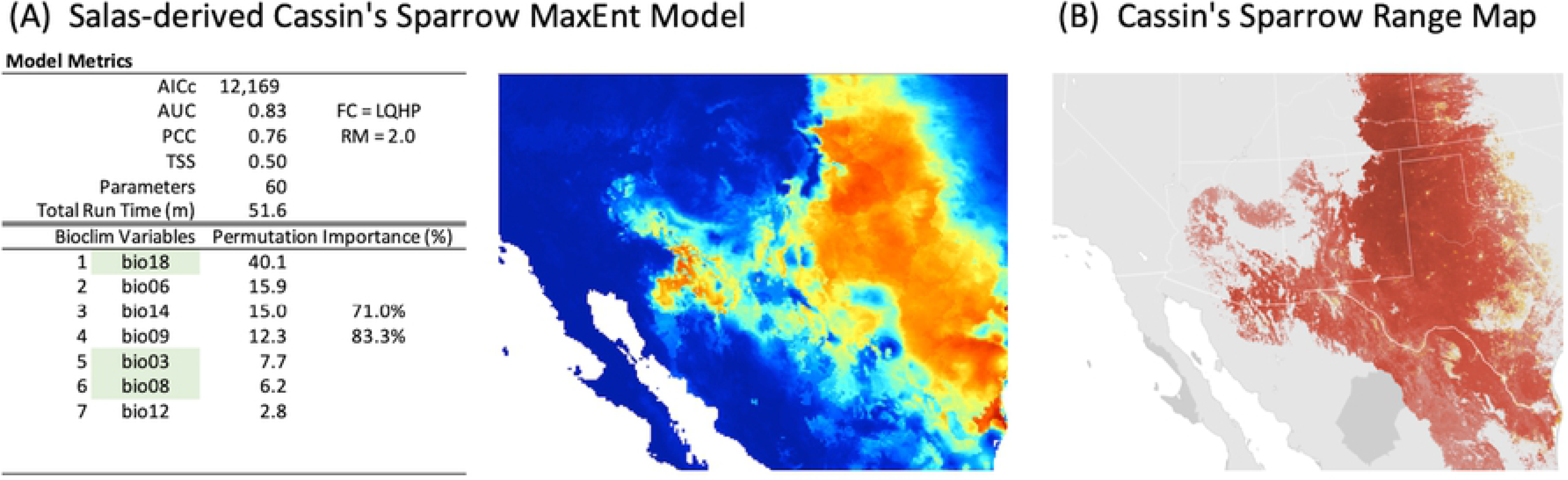
Cassin’s Sparrow baseline model and maps. Figure shows results from a MaxEnt run that builds on the Cassin’s Sparrow bioclimatic modeling work of Salas et al. [68] and reflects a more traditional approach to ENM (A) and Cassin’s Sparrow’s range map based on observational data (B). Highlighted variables indicate those that were also selected by MERRA/Max in the Bioclim use case. Range map provided by eBird (www.ebird.org), created 28 July 2020, and reprinted from [76] under a CC BY license, with permission from the Cornell Lab of Ornithology.

## Discussion

Climate change research is giving rise to new technology requirements at the intersection of big data, machine learning, and high-performance computing [77]. There are few places where this is more clearly seen than with studies focusing on the climate’s impact on species distribution and abundance [78]. For nearly twenty years, the ecological modeling community’s tool-of-choice for this work has been MaxEnt [16]. Few, if any, machine learning programs have been more widely used or more carefully studied [77,79–87]. Today, however, there is increasing interest in using GCM outputs as predictors in ENM [10–12], which has brought into focus one of the more challenging problems with existing machine learning systems: how to make them work with large, complex, feature-rich, high-dimensional data sets [88–94].

This study reflects our efforts to address this problem. Our approach to dimensionality reduction involves a parallel, out-of-core Monte Carlo selection method implemented in a high-performance, cloud computing setting. Monte Carlo optimizations are a way of finding approximate answers to problems that are solvable in principle but lack a practical means of solution [95]. Out-of-core (or external memory) algorithms process data sets that are too large to fit into a computer’s main memory [96,97]. They are currently a major focus of research in the machine learning community [97–99]. With MERRA/Max, we bring these concepts together to find a useful subset of predictors in a large collection of environmental variables in a reasonable amount of time. Early results are encouraging and suggest that the approach holds promise from both a technological and scientific perspective.

### Run-time performance and scaling properties

To begin, we have shown that MERRA/Max’s parallel implementation of the Monte Carlo selection algorithm scales linearly with respect to the average number of samples taken of each variable in the collection being screened. It can recruit additional processor cores to maintain a constant run time regardless of the size of the collection. In a best case, where sufficient processors are available for complete parallelism, MERRA/Max can screen a predictor data set of any size in the time it takes for a single MaxEnt run using only two predictors in the target collection. Collectively, these results confirm that near real-time performance and, in the vernacular of high-performance cloud computing, “infinite scalability” are achievable [100–102].

### Use case scenarios and selection behavior

MERRA/Max’s run-time performance and scaling properties are consistent with what one would expect of an “embarrassingly parallel” workload, where subtasks are completely independent and able to run concurrently. The more important question is: Does this approach actually benefit science? We try to assess MERRA/Max’s potential value to science by addressing three interrelated questions:

#### (1) Is MERRA/Max making useful, ecologically meaningful selections?

It is difficult to know how best to evaluate selection success in this work, given that we are proposing the Monte Carlo method as a preliminary screening step, which, presumably, would be followed by further refinements to the set of selected predictors based on the biology and ecology of the species, collinearity, or the other considerations that have traditionally guided ENM variable selection. Depending on the circumstances, post-selection refinement might mean additional winnowing, substitution or augmentation of the screened predictors with other variables, or that the selected variables are discarded altogether. That being said, the Bioclim use case seems to confirm that MERRA/Max’s selections are both valid and useful.

In the most general, qualitative sense, the habitat suitability map produced by the final model in the Bioclim use case is consistent with what is known about Cassin’s Sparrow’s range from observational records [76] (Fig 3A, Fig 4B). Likewise, the set of selected variables are consistent with what is known of the species’ natural history. Cassin’s Sparrow is a desert-adapted, ground-dwelling (and, notably, ground-nesting) species, whose breeding biology is exquisitely linked to conditions of temperature and precipitation and their consequent influence on vegetation availability, insect abundance, and terrestrial microclimates [43,47,49,52,103–105]. In fact, field studies over the past century suggest that Cassin’s Sparrow is an itinerant breeder, so responsive to temperature and precipitation that they make seasonal, inter-clutch moves within their range to find optimal conditions for breeding [43,104,106].

The Bioclim scenario’s ordered selection of bio18 (precipitation of the warmest quarter), bio03 (isothermality, i.e., temperature evenness, or how large the daily temperature variation is compared to its annual variation), bio05 (maximum temperature of the warmest month), bio13 (precipitation of the wettest month), and bio08 (mean temperature of wettest quarter) is entirely consistent with this picture. It is also largely consistent with the variables assembled by the Salas team using a more traditional approach to variable selection [68]. Both sets have bio18 and bio03 in common, both of these variables are highly influential, and both are known to be important determinants of range in arid-adapted birds, especially in desert and grassland species of conservation concern [68,78,107–113]. Where the two predictor data sets differ, for example, bio05 (maximum temperature of the warmest month) in the Bioclim use case vs. bio06 (minimum temperature of the coldest month) in the Salas-derived model, bio13 (precipitation of wettest month) vs. bio12 (annual precipitation) and bio 14 (precipitation of driest month), an argument could be made, in light of Cassin’s Sparrows distinctive seasonal breeding dynamics, which is likely influenced by the North American Monsoon, that MERRA/Max found the more relevant predictors [114–116].

From a quantitative perspective, the final model in MERRA/Max’s Bioclim scenario (Fig 3A) demonstrated strong evaluation metrics (AUC, PCC, TSS of 0.83, 0.75, 0.49 respectively) that compared favorably with those obtained in the Salas-derived model (0.83, 0.76, 0.50) (Fig 4A) [117]. A high degree of similarity between the Bioclim use case and Salas-derived model is further confirmed by their AICc values (12,232 and 12,169 respectively) and the results we obtained for the *D*, *I*, and *r* statistics (0.974, 0.999, and 0.997 respectively) [73]. The top four predictors in the Bioclim scenario accounted for 98% of overall permutation importance in the final model; the top four predictors in the Salas-derived model accounted for 83%.

While these results reflect an admittedly limited trial at this point, taken together, they suggest that MERRA/Max’s use in the Bioclim scenario produced a bioclimatic niche model for Cassin’s Sparrow that is ecologically reasonable, statistically robust, and at least as good (if not better) than what might be obtained in a traditional application of MaxEnt. This gives us confidence that the Monte Carlo method is, in fact, finding a useful subset of predictors in a larger pool of possible predictors.

#### (2) Does MERRA/Max create new research opportunities?

Another measure of MERRA/Max’s potential value to science is to consider whether new avenues of research are opened up with this technology. We believe they are, and the MERRA-2 use case helps explain why. Here we have a situation where the size and complexity of the target collection, as well as the obscure nature of the data, make *a priori* variable selection difficult. We have no immediate basis for distinguishing which variables in the MERRA-2 test collection might be the most important contributors to a final model. With 86 variables in play, even something as straightforward as correlation analysis provides little help, requiring nearly 3700 pair-wise comparisons in this case. Yet, going into this with no pre-vetting of the test collection whatsoever, we are struck by the ecological and biological relevance of the predictors selected by MERRA/Max in the MERRA-2 use case scenario.

In the same way that the Earth’s climate is ultimately driven by a balance between incoming and outgoing energy, so too is the natural history of Cassin’s Sparrow linked to the energetics of the species’ diurnal and seasonal activities and the locations where those activities occur [49]. Viewed this way, it makes ecological sense to see that MERRA/Max identified M39 (surface net downward shortwave flux assuming a clear day) as the most important variable for modeling Cassin’s Sparrow potential habitat in the MERRA-2 test collection. Likewise, a unique aspect of the Cassin’s Sparrow’s breeding biology is an energetically demanding skylark display in which males define and defend territories and secure mates by aerial flight songs. Wind has a pronounced impact on this behavior [49]. While its effectiveness as a proxy for surface conditions in this setting is unknown, it is notable that MERRA/Max identified a low-level zonal wind component, M03 (northward wind), as a top contributor. Finally, given Cassin’s Sparrow’s ground-dwelling habit and the importance of low-level environmental conditions to almost all aspects of the species’ life, it is not surprising, and also affirming, to see M05 (specific humidity), M04 (temperature), and M81 (bare soil evaporation) in MERRA/Max’s selection set.

Ultimately, the set of MERRA-2 variables selected by MERRA/Max seems to be a remarkably appropriate set of bioclimatic predictors for Cassin’s Sparrow. They are consistent with our mechanistic, process-based understanding of the bird’s natural history [49]. Further, in the use case scenario’s final MaxEnt model, we see a reasonable habitat suitability map based on our understanding of the species known range (Fig 4b), a relatively robust set of metrics (AUC 0.83, PCC 0.72, TSS 0.44), and the top four contributing variables accounting for a significant proportion (i.e., 97%) of overall permutation importance (Fig 3B). Taken together, it appears that MERRA/Max and the Monte Carlo selection method have detected a signal in the MERRA-2 data that has both biological and statistical significance for Cassin’s Sparrow [118,119].

What about the larger question regarding new research opportunities? A particular type of question that might be better addressed with MERRA/Max’s combination of data and technology concerns the conservation status of arid-adapted birds. Accurate assessments are often difficult with these species [78,107–113,120]. With Cassin’s Sparrow, for example, numerous studies over the past half-century have painted a confusing picture. Some find evidence for a retraction of viable habitat and declining regional populations [52], others find mixed results and too little data to establish with confidence an overall conservation status [47], and many sources identify the species as stable and of little immediate worry [121–124]. In recent work, of nine grassland birds of conservation concern, Cassin’s Sparrow was the only species to project gains in suitable habitat over the next fifty years [68]. This ambiguous picture is not unique to Cassin’s Sparrow; however, in this case, the species’ itinerant breeding habit no doubt contributes to the confusion: it is simply impossible to know what part of the bird’s population one is seeing at any given time.

Understanding the conservation status of a bird like Cassin’s Sparrow means being able to tease apart the species’ physiological capacity for seasonal response to weather from the species’ longer-term, adaptive response to a changing climate’s effect on the landscape [111]. Ultimately, one would like to distinguish short-term transformations in the bird’s bioclimatic niche within a coherent, long-running temporal framework. With four decades of climate attributes modeled on an hourly basis, reanalyses, such as MERRA-2, are uniquely able to provide the necessary high-temporal resolution, longitudinal environmental data space for this. Technology like MERRA/Max transforms MERRA-2 into a viable experimental sandbox. And, thanks to long-running citizen-scientist efforts, such as the U.S. Geological Survey’s North American Breeding Bird Survey (BBS) [125,126]; Cornell University’s eBird, Great Backyard Bird Count Surveys, and other projects [67,127,128]; Audubon’s Christmas Bird Counts [129,130]; and the wealth of online museum specimen records in resources like the Global Biodiversity Information Facility (GBIF) [131], the NSF-funded VertNet databases [132], and the U.S. Geological Survey’s BISON species occurrence database [133], there now exists widespread availability of avian observations that provide good coverage for the MERRA-2 time span, making multi-temporal scale investigations like this possible.

We close this section by considering briefly the most apparent technical shortcoming in the MERRA-2 use case: the inherently coarse spatial resolution of reanalysis data. Within the ecological research community, there has been a long-running clamor for higher resolution reanalysis products, to which various efforts are now responding [134–136]. We acknowledge this drawback, particularly with respect to final model construction. However, in the context of rapid prescreening, MERRA-2’s coarse resolution data may actually serve MERRA/Max’s intended purpose well. As we have seen, selection times are fast with coarse-resolution data, and any resolution shortcomings in the selected variables can be addressed in the refinement step, either by downscaling variables of interest or going to an alternative source for a higher resolution product, such as remote sensing data [137,138] or NatureServe’s high-resolution data sets [139]. At this point, however, we have no evidence that *screening* high-resolution data would return different recommendations than screening coarse-resolution data.

#### (3) Can MERRA/Max improve the ecological niche modeling process?

Finally, in evaluating potential benefits to science, we can consider whether a tool like MERRA/Max could improve the work practices of ecological modeling. Here again, we think it can. There is heightened awareness of the significance of dimensionality in understanding environmental spaces and the importance of variable selection in modeling those spaces [87,140,141]. This awareness is accompanied by a recognition that logistic difficulties often preclude examining large numbers of variables [81,142]. This has led to a search for alternative means of variable selection and calls for process automation [86,87,142–144]. A comprehensive review of these approaches is beyond the scope of this paper; however, it is worth noting that even among the most recent work in this area, many of the solutions put forward, such as manual prescreening for collinear variables, greater use of biological insight in variable selection, broader use of memory-resident machine learning-based analysis software, etc., do not, in general, scale well [145]. They are unlikely to accommodate the petabyte and even larger size data collections on the horizon.

Cobos et al. [86] provide a helpful framework for understanding where a technology like MERRA/Max could fit (Fig 5). The work of ENM can be thought of as a multi-step process ranging from initial data preparation and cleaning, to model calibration, final model construction, model evaluation, and the assessment of extrapolation risk. Among the tasks associated with data cleaning, the selection of viable predictors is crucial, time-consuming, and the place where a means for rapid, automatic, preselection could improve the overall workflow, especially if it enabled exploration of a large universe of predictors.

**Fig 5.**
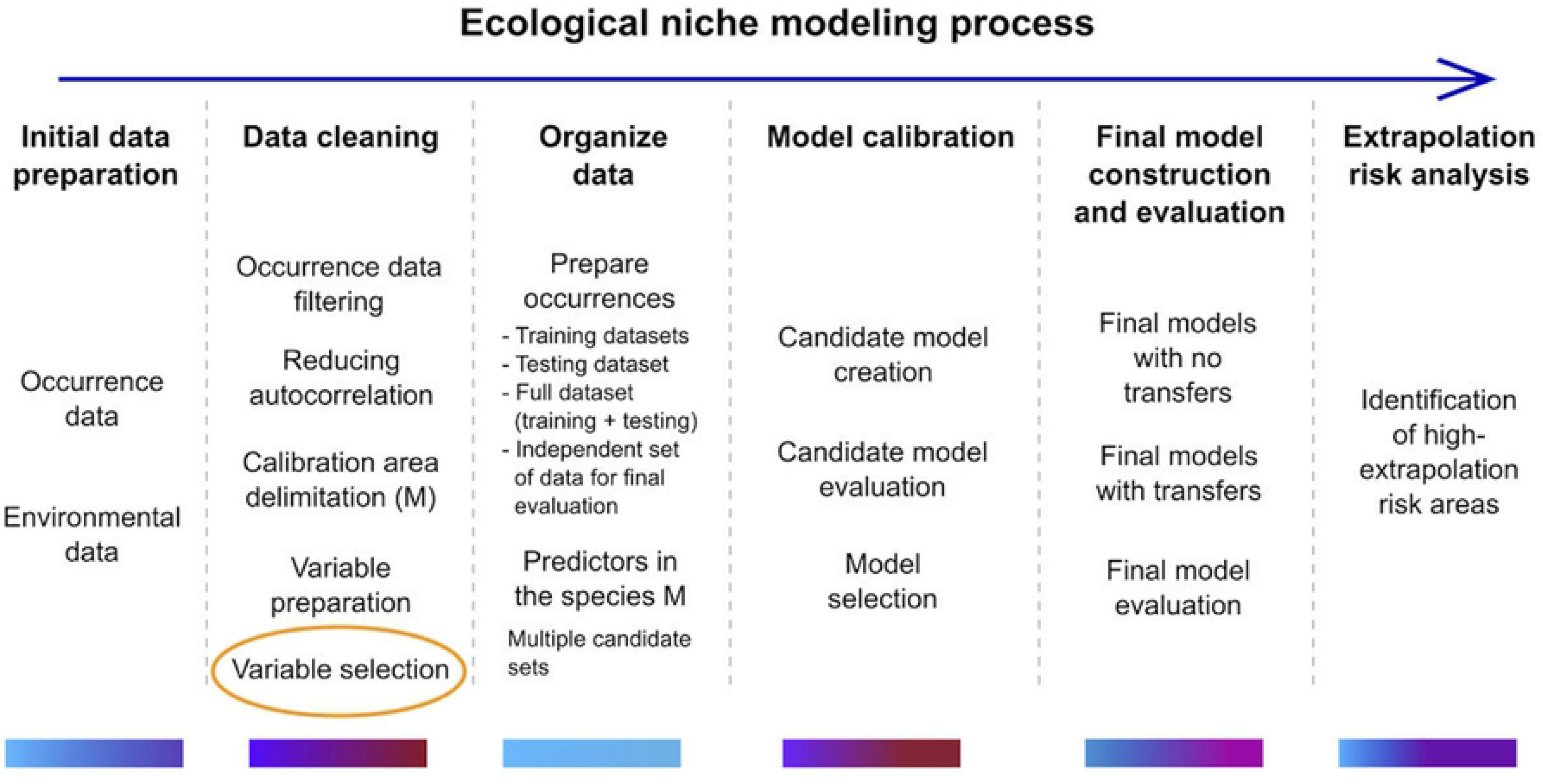
Ecological niche modeling (ENM) process. Schematic description of the ENM process. Color bars under each step reflect an approximate amount of time that may be needed, ranging from low (blue) to high (red). The use of MERRA/Max to prescreen a large collection of predictors could support variable selection in the data cleaning step. Image provided by [86] and adapted for use here under a CC-BY license.

### Future Work

Our plans for future work are being shaped by a vision where the technical complexities described here are abstracted away from the end-users, and low cost, easy access, and simplicity make MERRA/Max a practical and useful tool for the conservation research community. On the technology front, our next steps will focus on developing a “cloud bursting” capability that will allow MERRA/Max ensembles to migrate from NASA’s private ADAPT science cloud to a public commercial cloud in response to resource demands that outstrip local capacity. This will allow us to scale MERRA/Max to larger data sets and more demanding science questions. For example, we would like to make better use of our experimental MERRA-2 test collection, which spans 40 years and includes monthly and weekly maximum, minimum, and average values for each of the collection’s 86 variables, a total of N = 660,480 files. Beyond that, expanding MERRA/Max to accommodate all of MERRA-2’s 600-plus variables is a more challenging long-term goal that could open interesting new research opportunities for the ENM community. Other technical improvements on the horizon include the use of refined selection criteria and automatic stopping rules that would enable true convergence in the selection process. Finally, given that each step in our use case scenarios is carried out by a program that could readily participate in an orchestrated workflow, we would like to know whether a practical level of automation for the entire MaxEnt-enabled ENM process might be possible.

On the science front, we plan to pursue two distinct threads of development. First, we want to follow up on the example posed in the previous section and see if MERRA/Max and the MERRA-2 test collection could provide a better understanding of the true conservation status of Cassin’s Sparrow by evaluating the species’ bioclimatic niche at multiple time scales over the past four decades. Pursuing that question for Cassin’s Sparrow alongside other grassland birds of conservation concern, such as those studied by Salas et al. [68], would provide the additional benefit of extending our experiences to other species and would allow us to test the ecological and conservation applicability of this technology, which we view as an important next step.

The second thread of science development will examine the extensibility of this approach to other types of problems. For example, we have used MERRA/Max and MaxEnt to study the hydrological cycle in NASA’s Arctic-Boreal Vulnerability Experiment (ABoVE) [146]. Changes to the hydrological cycle in the Arctic are particularly complex, because observed and projected warming directly impacts permafrost and leads to variable responses in surface water extent [147]. In preliminary work, using the locations of observed increases and decreases in surface water extent as dependent variables (in essence, treating them as “pseudo” species occurrences), the technologies and techniques described here successfully replicated observed patterns of surface water change [148]. If these findings are validated by further experimentation, the view of how MERRA/Max, the MERRA-2 reanalysis, and MaxEnt might be applied to studies of climate change and its impact on the Earth system becomes significantly broadened.

## Conclusions

In this paper, we have described a prototype system called MERRA/Max that implements a feature selection approach to dimensionality reduction that is specifically intended to enable direct use of GCM outputs in ENM. The system accomplishes this reduction through a Monte Carlo optimization in which many independent MaxEnt runs operating on a species occurrence file and a small set of variables randomly selected from a large collection of variables converges on an estimate of the top contributing predictors in the larger collection. These top predictors become candidates for consideration in the variable selection step of the ENM process. MERRA/Max’s Monte Carlo algorithm operates on files stored in the underlying filesystem and is thus scalable to large data sets. We implemented its program components using commercial, off-the-shelf software. These components can run independently as parallel processes in a high-performance cloud computing environment to yield near real-time performance.

Within this framework, variable selection is guided by the indirect biological influences injected into MERRA/Max’s variable reduction process by the species occurrence files. We find evidence for this tailoring of results in our use case scenarios. In preliminary tests using a single bird species and observations from a single year, MERRA/Max selected biologically valid and familiar climatological predictors from the classic Bioclim collection of environmental variables. MERRA/Max also selected biologically and ecologically meaningful predictors from a larger and much more diverse set of environmental variables derived from NASA’s MERRA-2 reanalysis. Our experience is limited at this point, but we feel that these results point to a technological approach that could expand the use of GCM outputs in ENM, streamline the modeling process, and, eventually, enable automated bioclimatic modeling as a cloud service.

## Supporting Information

**S1 Appendix. MERRA/Max Parameterization.** Overview of MERRA/Max’s default screening parameters and supporting documentation.

**S2 File. Data and Scripts.** Compressed file folder containing input data and example scripts used in the study.

## Acknowledgements

We thank Glenn Tamkin, Roger Gill, Jian Li, Savannah Strong, Tom Maxwell, Mary Aronne, and Caleb Spradlin for their many contributions and ongoing technical support of these efforts. Dr. Virginia Seamster of the New Mexico Department of Game and Fish provided valuable advice and much-appreciated editorial input on an earlier draft of the manuscript. Resources supporting this work were provided by the NASA High-End Computing (HEC) Program through the NASA Center for Climate Simulation (NCCS) at Goddard Space Flight Center.

## References

1. Peterson AT, Soberón J, Pearson RG, Anderson RP, Martínez-Meyer E, Nakamura M, et al. Ecological niches and geographic distributions (MPB-49). Princeton University Press; 2011.

2. Schmidt-Lebuhn AN, Knerr NJ, Miller JT, Mishler BD. Phylogenetic diversity and endemism of Australian daisies (Asteraceae). Journal of Biogeography. 2015;42: 1114–1122. doi:10.1111/jbi.12488

3. Cardoso-Leite R, Vilarinho AC, Novaes MC, Tonetto AF, Vilardi GC, Guillermo-Ferreira R. Recent and future environmental suitability to dengue fever in Brazil using species distribution model. Transactions of The Royal Society of Tropical Medicine and Hygiene. 2014;108: 99–104. doi:10.1093/trstmh/trt115

4. Franklin J. Species distribution models in conservation biogeography: developments and challenges. Diversity Distrib. 2013;19: 1217–1223. doi:10.1111/ddi.12125

5. Jiménez-Valverde A, Peterson AT, Soberón J, Overton JM, Aragón P, Lobo JM. Use of niche models in invasive species risk assessments. Biol Invasions. 2011;13: 2785–2797. doi:10.1007/s10530-011-9963-4

6. Muttaqin LA, Murti SH, Susilo B. MaxEnt (Maximum Entropy) model for predicting prehistoric cave sites in Karst area of Gunung Sewu, Gunung Kidul, Yogyakarta. In: Wibowo SB, Rimba AB, A. Aziz A, Phinn S, Sri Sumantyo JT, Widyasamratri H, et al., editors. Sixth Geoinformation Science Symposium. Yogyakarta, Indonesia: SPIE; 2019. p. 3. doi:10.1117/12.2543522

7. Searcy CA, Shaffer HB. Do Ecological Niche Models Accurately Identify Climatic Determinants of Species Ranges? The American Naturalist. 2016;187: 423–435. doi:10.1086/685387

8. Harris RMB, Grose MR, Lee G, Bindoff NL, Porfirio LL, Fox-Hughes P. Climate projections for ecologists: Climate projections for ecologists. Wiley Interdisciplinary Reviews: Climate Change. 2014;5: 621–637. doi:10.1002/wcc.291

9. Edwards PN. A Vast Machine: Computer Models, Climate Data, and the Politics of Global Warming. Cambridge, MA: MIT Press; 2010.

10. Cavanagh RD, Murphy EJ, Bracegirdle TJ, Turner J, Knowland CA, Corney SP, et al. A Synergistic Approach for Evaluating Climate Model Output for Ecological Applications. Frontiers in Marine Science. 2017;4: 308. doi:10.3389/fmars.2017.00308

11. Radosavljevic A, Anderson RP. Making better MaxEnt models of species distributions: complexity, overfitting and evaluation. Araújo M, editor. Journal of Biogeography. 2014;41: 629–643. doi:10.1111/jbi.12227

12. Heinze G, Wallisch C, Dunkler D. Variable selection – A review and recommendations for the practicing statistician. Biometrical Journal. 2018;60: 431–449. doi:10.1002/bimj.201700067

13. Schnase JL, Carroll ML, Gill RL, Tamkin GS, Li J, Strong SL, et al. Toward a Monte Carlo approach to selecting climate variables in MaxEnt. PLOS ONE. 2021;16: e0237208. doi:10.1371/journal.pone.0237208

14. Elith J, Phillips SJ, Hastie T, Dudík M, Chee YE, Yates CJ. A statistical explanation of MaxEnt for ecologists. Diversity and distributions. 2011;17: 43–57.

15. Phillips SJ, Anderson RP, Dudík M, Schapire RE, Blair ME. Opening the black box: An open-source release of Maxent. Ecography. 2017;40: 887–893.

16. Phillips SJ. A Brief Tutorial on Maxent. AT&T Research. 2005;190: 231–259.

17. Schnase JL, Smith JA, Stohlgren TJ, Graves S, Trees C. Biological Invasions: a Challenge in Ecological Forecasting. IEEE International Geoscience and Remote Sensing Symposium. 2002. doi:10.1109/igarss.2002.1024961.

18. Beauchamp VB, Koontz SM, Suss C, Hawkins C, Kyde KL, Schnase JL. An Introduction to Oplismenus Undulatifolius (Ard.) Roem. & Schult (Wavyleaf Basketgrass), a Recent Invader in Mid-Atlantic Forest Understories 1,2. The Journal of the Torrey Botanical Society. 2013;140: 391–413. doi:10.3159/torrey-d-13-00033.1.

19. Morisette JT, Jarnevich CS, Ullah A, Cai W, Pedelty JA, Gentle JE, et al. A Tamarisk Habitat Suitability Map for the Continental United States. Frontiers in Ecology and the Environment. 2006;4: 11–17. doi:10.1890/1540-9295(2006)004

20. Schnase JL, Carroll ML, Weber KT, Brown ME, Gill RL, Wooten M, et al. RECOVER: An Automated, Cloud-Based Decision Support System for Post-Fire Rehabilitation Planning. ISPRS - International Archives of the Photogrammetry, Remote Sensing and Spatial Information Sciences XL-1. 2014. pp. 363–70. doi:10.5194/isprsarchives-xl-1-363-2014

21. Schnase JL, Most N, Gill R, Ma P. The Invasive Species Forecasting System. 2009 17th International Conference on Geoinformatics. 2009. doi:10.1109/geoinformatics.2009.5293333.

22. Hu F, Yang C, Schnase JL, Duffy DQ, Xu M, Bowen MK, et al. ClimateSpark: An in-memory distributed computing framework for big climate data analytics. Computers & Geosciences. 2018;115: 154–166. doi:10.1016/j.cageo.2018.03.011

23. Schnase JL, Lee TJ, Mattmann CA, Lynnes CS, Cinquini L, P. M. Ramirez, et al. Big Data Challenges in Climate Science: Improving the next-generation cyberinfrastructure. IEEE Geoscience and Remote Sensing Magazine. 2016;4: 10–22. doi:10.1109/MGRS.2015.2514192

24. Schnase JL. Climate Analytics as a Service. Cloud Computing in Ocean and Atmospheric Sciences. 2016. pp. 187–219. doi:10.1016/b978-0-12-803192-6.00011-6

25. Schnase JL, Cushing J, Frame M, Frondorf A, Landis E, Maier D, et al. Information technology challenges of biodiversity and ecosystems informatics. Information Systems. 2003;28: 339–345. doi:10.1016/S0306-4379(02)00070-4

26. Schnase JL, Duffy DQ, McInerney MA, Webster WP, Lee TJ. Climate Analytics as a Service. Proceedings of the 2014 Conference on Big Data from Space (BiDS. Frascati: European Space Agency (ESA; 2014. pp. 90–94. doi:10.2788/1823.

27. Schnase JL, Duffy DQ, Tamkin GS, Nadeau D, Thompson JH, Grieg CM, et al. MERRA Analytic Services: Meeting the Big Data Challenges of Climate Science through Cloud-Enabled Climate Analytics-as-a-Service. Computers, Environment and Urban Systems. 2017;61: 198–211. doi:10.1016/j.compenvurbsys.2013.12.003.

28. Carriere L, Potter GL, Hertz J, Peters J, Maxwell TP, Strong S, et al. CREATE-IP and CREATE-V: Data and Services Update. AGU Fall Meeting Abstracts. 2017. pp. IN21D–0064.

29. Cinquini L, Crichton D, Mattmann C, Harney J, Shipman G, Wang F, et al. The Earth System Grid Federation: An open infrastructure for access to distributed geospatial data. Future Generation Computer Systems. 2014;36: 400–417.

30. Maxwell TP, Potter GL, Carriere L, Duffy D. The Earth Data Analytic Services Framework. AGU Fall Meeting Abstracts. 2019. pp. IN13B–0720.

31. Tamkin G, Schnase JL, Duffy D, Li J, Strong S, Thompson JH. The NASA Reanalysis Ensemble Service-Advanced Capabilities for Integrated Reanalysis Access and Intercomparison. AGU Fall Meeting Abstracts. 2017. pp. IN21D–0065.

32. GES DISC - Goddard Earth Science Data and Information Services Center. [cited 26 May 2021]. Available: https://disc.gsfc.nasa.gov/

33. NASA Case Study – Amazon Web Services (AWS). In: Amazon Web Services, Inc. [Internet]. 2021 [cited 26 May 2021]. Available: https://aws.amazon.com/partners/success/nasa-image-library/

34. Research and Technical Computing on Amazon Web Services (AWS). In: Amazon Web Services, Inc. [Internet]. 2021 [cited 26 May 2021]. Available: https://aws.amazon.com/government-education/research-and-technical-computing/

35. Google Cloud offers global support for academic research. In: Google [Internet]. 2019 [cited 26 May 2021]. Available: https://blog.google/products/google-cloud/google-cloud-offers-global-support-for-academic-research/

36. Our head’s in the cloud, but we’re keeping the earth in mind. In: Google Cloud Blog [Internet]. 2019 [cited 26 May 2021]. Available: https://cloud.google.com/blog/topics/google-cloud-next/our-heads-in-the-cloud-but-were-keeping-the-earth-in-mind/

37. Cloud Computing Services | Microsoft Azure. 2021 [cited 26 May 2021]. Available: https://azure.microsoft.com/en-us/

38. Data Science Virtual Machines | Microsoft Azure. 2021 [cited 26 May 2021]. Available: https://azure.microsoft.com/en-us/services/virtual-machines/data-science-virtual-machines/

39. ADAPT. In: ADAPT | NASA Center for Climate Simulation [Internet]. [cited 15 Mar 2021]. Available: https://www.nccs.nasa.gov/systems/ADAPT

40. R: The R Project for Statistical Computing. [cited 22 May 2020]. Available: https://www.r-project.org/

41. Muscarella R, Galante PJ, Soley-Guardia M, Boria RA, Kass JM, Anderson MU and RP. ENMeval: Automated Runs and Evaluations of Ecological Niche Models. 2020. Available: https://CRAN.R-project.org/package=ENMeval

42. Maxent Version 3.4.1 Download Site. In: Maxent Version 3.4.1 Download Site [Internet]. [cited 22 May 2020]. Available: https://biodiversityinformatics.amnh.org/open_source/maxent/

43. Dunning, Jr. JB, Bowers, Jr. RK, Suter SJ, Bock CE. Cassin’s Sparrow (Peucaea cassinii), Version 1.0. In: Birds of the World (P. G. Rodewald, Editor) [Internet]. 2020 [cited 22 May 2020]. Available: https://doi.org/10.2173/bow.casspa.01

44. Fick SE, Hijmans RJ. WorldClim 2: new 1-km spatial resolution climate surfaces for global land areas. International Journal of Climatology. 2017;37: 4302–4315. doi:10.1002/joc.5086

45. Worldclim bioclimatic variables. 2020 [cited 22 May 2020]. Available: https://worldclim.org/data/worldclim21.html

46. GBIF.org (21 February 2019) GBIF Occurrence Download https://doi.org/10.15468/dl.0s8yak.

47. Ruth JM. Cassin’s Sparrow Status Assessment and Conservation Plan. Biological Technical Publication BTP-R6002-2000. Denver, CO: U.S. Department of the Interior, Fish and Wildlife Service; 2000.

48. Schnase JL, Maxwell TC. Use of song patterns to identify individual male Cassin’s Sparrows. Journal of Field Ornithology. 1989;60: 12–19.

49. Schnase JL, Grant WE, Maxwell TC, Leggett JJ. Time and energy budgets of Cassin’s sparrow (Aimophila cassinii) during the breeding season: evaluation through modelling. Ecological Modelling. 1991;55: 285–319.

50. GDAL/OGR Geospatial Data Abstraction Software Library. Open Source Geospatial Foundation; 2020. Available: https://gdal.org/

51. Hijmans RJ, Phillips S, Elith J, Leathwick J. dismo: Species Distribution Modeling. 2017. Available: https://CRAN.R-project.org/package=dismo

52. Lynn J. Cassin’s Sparrow (Aimophila cassinii): A Technical Conservation Assessment. USDA Forest Service, Species Conservation Project. Rocky Mountain Region. 2006;46.

53. Pradhan P. Strengthening MaxEnt modelling through screening of redundant explanatory bioclimatic variables with variance inflation factor analysis. Researcher. 2016;8: 29–34.

54. Warren DL, Glor RE, Turelli M. ENMTools: a toolbox for comparative studies of environmental niche models. Ecography. 2010 [cited 27 Mar 2020]. doi:10.1111/j.1600-0587.2009.06142.x

55. Muscarella R, Galante PJ, Soley-Guardia M, Boria RA, Kass JM, Uriarte M, et al. ENMeval: An R package for conducting spatially independent evaluations and estimating optimal model complexity for Maxent ecological niche models. Methods in Ecology and Evolution. 2014;5: 1198–1205. doi:10.1111/2041-210X.12261

56. H. Akaike. A new look at the statistical model identification. IEEE Transactions on Automatic Control. 1974;19: 716–723. doi:10.1109/TAC.1974.1100705

57. IPCC — Intergovernmental Panel on Climate Change. 2020 [cited 14 Mar 2020]. Available: https://www.ipcc.ch/

58. Gelaro R, Mccarty W, Su MJ, Todling R, Molod A, Takacs L, et al. The Modern-Era Retrospective Analysis for Research and Applications, Version 2 (MERRA-2). JOURNAL OF CLIMATE. 2017;30: 36.

59. Bosilovich MG, Lucchesi R, Suarez M. MERRA-2: File Specification. GMAO Office Note. 2016;9: 1–73.

60. MERRA-2. 2020 [cited 19 Mar 2020]. Available: https://gmao.gsfc.nasa.gov/reanalysis/MERRA-2/

61. Beaumont LJ, Hughes L, Poulsen M. Predicting species distributions: use of climatic parameters in BIOCLIM and its impact on predictions of species’ current and future distributions. Ecological Modelling. 2005;186: 251–270. doi:10.1016/j.ecolmodel.2005.01.030

62. Booth TH, Nix HA, Busby JR, Hutchinson MF. BIOCLIM: the first species distribution modelling package, its early applications and relevance to most current MaxEnt studies. Franklin J, editor. Diversity Distrib. 14AD;20: 1–9. doi:10.1111/ddi.12144

63. Hijmans RJ, Cameron SE, Parra JL, Jones PG, Jarvis A. Very high resolution interpolated climate surfaces for global land areas. International Journal of Climatology. 2005;25: 1965–1978. doi:10.1002/joc.1276

64. O’Donnell MS, Ignizio D a. Bioclimatic Predictors for Supporting Ecological Applications in the Conterminous United States. Reston, VA: US Geological Survey; 2012 p. 10. Report No.: 691. Available: https://pubs.usgs.gov/ds/691/

65. Vega GC, Pertierra LR, Olalla-Tárraga MÁ. Data from: MERRAclim, a high-resolution global dataset of remotely sensed bioclimatic variables for ecological modelling. Dryad; 2018. p. 8324359732 bytes. doi:10.5061/DRYAD.S2V81

66. Hoyer S, Hamman JJ. xarray: N-D labeled Arrays and Datasets in Python. Journal of Open Research Software. 2017;5: 10. doi:10.5334/jors.148

67. (Peucaea cassinii) - Species Map - eBird. 2020 [cited 31 May 2020]. Available: https://ebird.org/map/casspa

68. Salas EAL, Seamster VA, Boykin KG, Harings NM, Dixon KW, Department of Fish, Wildlife and Conservation Ecology, New Mexico State University, Las Cruces, New Mexico 88003, USA. Modeling the impacts of climate change on Species of Concern (birds) in South Central U.S. based on bioclimatic variables. AIMS Environmental Science. 2017;4: 358–385. doi:10.3934/environsci.2017.2.358

69. Fielding AH, Bell JF. A review of methods for the assessment of prediction errors in conservation presence/absence models. Environmental Conservation. 1997;24: 38–49. doi:10.1017/S0376892997000088

70. Warren DL, Seifert SN. Ecological niche modeling in Maxent: the importance of model complexity and the performance of model selection criteria. Ecological Applications. 2011;21: 335–342. doi:10.1890/10-1171.1

71. Allouche O, Tsoar A, Kadmon R. Assessing the accuracy of species distribution models: prevalence, kappa and the true skill statistic (TSS): Assessing the accuracy of distribution models. Journal of Applied Ecology. 2006;43: 1223–1232. doi:10.1111/j.1365-2664.2006.01214.x

72. Frans VF. True Skill Statistic (TSS) Calculation across Multiple Maxent Runs. Michigan State University; 2018 p. 5.

73. Warren DL, Glor RE, Turelli M. Environmental niche equivalency versus conservatism: quantitative approaches to niche evolution. Evolution. 2008;62: 2868–2883. doi:10.1111/j.1558-5646.2008.00482.x

74. Schoener TW. The Anolis Lizards of Bimini: Resource Partitioning in a Complex Fauna. Ecology. 1968;49: 704–726. doi:10.2307/1935534

75. Rodgers JL, Nicewander WA. Thirteen Ways to Look at the Correlation Coefficient. The American Statistician. 1988;42: 59–66. doi:10.2307/2685263

76. Fink D, Auer T, Johnston A, Strimas-Mackey M, Robinson O, Ligocki S, et al. Cassin’s Sparrow - Abundance map - eBird Status and Trends. In: eBird Status and Trends, Data Version: 2018; Released: 2020 [Internet]. 2020 [cited 5 Oct 2020]. Available: https://ebird.org/ebird/science/status-and-trends/casspa/abundance-map

77. Araújo M, New M. Ensemble forecasting of species distributions. Trends in Ecology & Evolution. 2007;22: 42–47. doi:10.1016/j.tree.2006.09.010

78. Huntley B, Barnard P, Altwegg R, Chambers L, Coetzee BWT, Gibson L, et al. Beyond bioclimatic envelopes: dynamic species’ range and abundance modelling in the context of climatic change. Ecography. 2010 [cited 13 Mar 2020]. doi:10.1111/j.1600-0587.2009.06023.x

79. Feng X, Park DS, Walker C, Peterson AT, Merow C, Papeş M. A checklist for maximizing reproducibility of ecological niche models. Nature Ecology & Evolution. 2019;3: 1382–1395. doi:10.1038/s41559-019-0972-5

80. Morales NS, Fernández IC, Baca-González V. MaxEnt’s parameter configuration and small samples: are we paying attention to recommendations? A systematic review. PeerJ. 2017;5: e3093. doi:10.7717/peerj.3093

81. Zeng Y, Low BW, Yeo DCJ. Novel methods to select environmental variables in MaxEnt: A case study using invasive crayfish. Ecological Modelling. 2016;341: 5–13. doi:https://doi.org/10.1016/j.ecolmodel.2016.09.019

82. Araújo MB, Anderson RP, Barbosa AM, Beale CM, Dormann CF, Early R, et al. Standards for distribution models in biodiversity assessments. Science Advances. 2019;5: eaat4858.

83. Qiao H, Soberón J, Peterson AT. No silver bullets in correlative ecological niche modelling: insights from testing among many potential algorithms for niche estimation. Kriticos D, editor. Methods in Ecology and Evolution. 2015;6: 1126–1136. doi:10.1111/2041-210X.12397

84. Ashraf U, Peterson AT, Chaudhry MN, Ashraf I, Saqib Z, Rashid Ahmad S, et al. Ecological niche model comparison under different climate scenarios: a case study of *Olea* spp. in Asia. Ecosphere. 2017;8: e01825. doi:10.1002/ecs2.1825

85. Guisan A, Zimmermann NE. Predictive habitat distribution models in ecology. Ecological modelling. 2000;135: 147–186.

86. Cobos ME, Peterson AT, Barve N, Osorio-Olvera L. kuenm: an R package for detailed development of ecological niche models using Maxent. PeerJ. 2019;7: e6281. doi:10.7717/peerj.6281

87. Cobos ME, Peterson AT, Osorio-Olvera L, Jiménez-García D. An exhaustive analysis of heuristic methods for variable selection in ecological niche modeling and species distribution modeling. Ecological Informatics. 2019;53: 100983. doi:10.1016/j.ecoinf.2019.100983

88. Qiu J, Wu Q, Ding G, Xu Y, Feng S. A survey of machine learning for big data processing. EURASIP J Adv Signal Process. 2016;2016: 67. doi:10.1186/s13634-016-0355-x

89. Obermeyer Z, Emanuel EJ. Predicting the Future - Big Data, Machine Learning, and Clinical Medicine. N Engl J Med. 2016;375: 1216–1219. doi:10.1056/NEJMp1606181

90. Bailly S, Meyfroidt G, Timsit J-F. What’s new in ICU in 2050: big data and machine learning. Intensive Care Medicine. 2018;44: 1524–1527. doi:10.1007/s00134-017-5034-3

91. van der Maaten L, Postma E, van den Herik J. Dimensionality Reduction: A Comparative Review. The Netherlands: Tilburg University; 2009 p. 36. Report No.: TiCC TR 2009-005. Available: https://members.loria.fr/moberger/Enseignement/AVR/Exposes/TR_Dimensiereductie.pdf

92. Guyon I, Elisseeff A. An Introduction to Variable and Feature Selection. Journal of Machine Learning Research. 2003;3: 1157–1182.

93. M. Espadoto, R. M. Martins, A. Kerren, N. S. T. Hirata, A. C. Telea. Towards a Quantitative Survey of Dimension Reduction Techniques. IEEE Transactions on Visualization and Computer Graphics. 2019; 1–1. doi:10.1109/TVCG.2019.2944182

94. Feng X, Park DS, Liang Y, Pandey R, Papeş M. Collinearity in ecological niche modeling: Confusions and challenges. Ecology and Evolution. 2019;9: 10365–10376. doi:10.1002/ece3.5555

95. Kroese DP, Brereton T, Taimre T, Botev ZI. Why the Monte Carlo method is so important today: Why the MCM is so important today. Wiley Interdisciplinary Reviews: Computational Statistics. 2014;6: 386–392. doi:10.1002/wics.1314

96. Ito Y, Imai H, Duc TL, Negishi Y, Kawachiya K, Matsumiya R, et al. Profiling based Out-of-core Hybrid Method for Large Neural Networks. arXiv:190705013 [cs]. 2019 [cited 11 Jan 2021]. Available: http://arxiv.org/abs/1907.05013

97. Chen T, Xu B, Zhang C, Guestrin C. Training Deep Nets with Sublinear Memory Cost. arXiv:160406174 [cs]. 2016 [cited 11 Jan 2021]. Available: http://arxiv.org/abs/1604.06174

98. Hanlon J. How To Solve The Memory Challenges Of Deep Neural Networks. In: TOPBOTS [Internet]. 2017 [cited 13 May 2021]. Available: https://www.topbots.com/how-solve-memory-challenges-deep-learning-neural-networks-graphcore/

99. Zhang B. A Solution to the Memory Limit Challenge in Big Data Machine Learning. In: Medium [Internet]. 2018 [cited 13 May 2021]. Available: https://petuum.medium.com/a-solution-to-the-memory-limit-challenge-in-big-data-machine-learning-49783a72088b

100. Bicer T, Chiu D, Agrawal G. A Framework for Data-Intensive Computing with Cloud Bursting. 2011 IEEE International Conference on Cluster Computing. 2011. pp. 169–177. doi:10.1109/CLUSTER.2011.21

101. Pham B, Jones RC, Shalaan M. Analysis of Cloud Bursting from the Openstack Infrastructure to AWS. 2020 IEEE Cloud Summit. 2020. pp. 114–118. doi:10.1109/IEEECloudSummit48914.2020.00037

102. Guo T, Sharma U, Shenoy P, Wood T, Sahu S. Cost-Aware Cloud Bursting for Enterprise Applications. ACM Trans Internet Technol. 2014;13. doi:10.1145/2602571

103. Oberholser HC. Cassin’s Sparrow, “Aimophila cassinii” (Woodhouse). 1st ed. In: Kincaid, Jr. EB, editor. Bird Life of Texas. 1st ed. Austin: University of Texas Press; 1974. pp. 920–921.

104. Williams FC, LeSassier AL. Cassin’s Sparrow. In: Austin OL, editor. Life Histories of North American Cardinals, Grosbeaks, Buntings, Towhees, Finches, Sparrows, and Allies, Order Passeriformes, Family Fringillidae: (in 3vols) Part 2, Genera Pipilo (part) Through Spizella. New York: Dover; 1968. pp. 981–990.

105. Woodhouse SW. Zonotrichia Cassinii, nobis. Proceedings of the Academy of Natural Science of Philadelphia. Philadelphia: Merrihow and Thompson; 1852. pp. 60–61. Available: https://www.biodiversitylibrary.org/item/17888#page/7/mode/1up

106. Ohmart RD. Dual breeding ranges in Cassin’s sparrow (Aimophila cassinii). In: Hoff CC, Riedesel ML, editors. Physiological systems in semiarid environments. Albuquerque, NM: University of New Mexico Press; 1969. p. 105.

107. Norman JA, Christidis L. Ecological opportunity and the evolution of habitat preferences in an arid-zone bird: implications for speciation in a climate-modified landscape. Scientific Reports. 2016;6: 19613. doi:10.1038/srep19613

108. Hubbard JP. Avian evolution in the aridlands of North America. The Living Bird. 1974; 155–196.

109. Sohl TL. The Relative Impacts of Climate and Land-Use Change on Conterminous United States Bird Species from 2001 to 2075. Romanach SS, editor. PLoS ONE. 2014;9: e112251. doi:10.1371/journal.pone.0112251

110. Rosenberg KV, Dokter AM, Blancher PJ, Sauer JR, Smith AC, Smith PA, et al. Decline of the North American avifauna. Science. 2019;366: 120. doi:10.1126/science.aaw1313

111. Reside AE, VanDerWal JJ, Kutt AS, Perkins GC. Weather, Not Climate, Defines Distributions of Vagile Bird Species. Hector A, editor. PLoS ONE. 2010;5: e13569. doi:10.1371/journal.pone.0013569

112. Iknayan KJ, Beissinger SR. Collapse of a desert bird community over the past century driven by climate change. Proc Natl Acad Sci USA. 2018;115: 8597. doi:10.1073/pnas.1805123115

113. Heenan CB, Seymour RS. The Effect of Wind on the Rate of Heat Loss from Avian Cup-Shaped Nests. Brigham RM, editor. PLoS ONE. 2012;7: e32252. doi:10.1371/journal.pone.0032252

114. Liebmann B. Characteristics of North American Summertime Rainfall with Emphasis on the Monsoon. American Meteorological Society Jounal of Climate. 2008;21: 1277–1294.

115. NOAA. The North American Monsoon. NOAA NWS Climate Prediction Center; 2019 p. 25. Available: https://www.cpc.ncep.noaa.gov/products/outreach/Report-to-the-Nation-Monsoon_aug04.pdf

116. Hansen H. Skylarking Cassin’s Sparrows in Southeast Arizona. In: ABA Blog [Internet]. 2019 [cited 23 Jun 2021]. Available: https://blog.aba.org/2019/08/skylarking-cassins-sparrows-in-southeast-arizona.html

117. West AM, Kumar S, Brown CS, Stohlgren TJ, Bromberg J. Field validation of an invasive species Maxent model. Ecological Informatics. 2016;36: 126–134. doi:10.1016/j.ecoinf.2016.11.001

118. Cuddington K, Fortin M-J, Gerber LR, Hastings A, Liebhold A, O’Connor M, et al. Process-based models are required to manage ecological systems in a changing world. Ecosphere. 2013;4: 1–12. doi:10.1890/ES12-00178.1

119. Kendall BE, Briggs CJ, Murdoch WW, Turchin P, Ellner SP, McCauley E, et al. Why do populations cycle? A synthesis of statistical and mechanistic modeling approaches. 1999;80: 17.

120. Lipschutz ML. Effects of Drought and Grazzing on Land Bird Populations in South Texas. MS, Range and Wildlife Management, Texas A&M University-Kingsville. 2016.

121. Cassin’s Sparrow - Whatbird.com. [cited 22 May 2021]. Available: https://identify.whatbird.com/obj/278/overview/cassins_sparrow.aspx

122. Cassin’s Sparrow. In: Audubon [Internet]. 2014 [cited 22 May 2021]. Available: https://www.audubon.org/field-guide/bird/cassins-sparrow

123. Cassin’s Sparrow (Peucaea cassinii) - BirdLife species factsheet. [cited 22 May 2021]. Available: http://datazone.birdlife.org/species/factsheet/22721272

124. Cassin’s Sparrow Life History, All About Birds, Cornell Lab of Ornithology. [cited 22 May 2021]. Available: https://www.allaboutbirds.org/guide/Cassins_Sparrow/lifehistory

125. Sauer JR, Link WA, Fallon JE, Pardieck KL, Ziolkowski DJ. The North American Breeding Bird Survey 1966-2011: Summary Analysis and Species Accounts. North American Fauna. 2013;79: 1–32. doi:10.3996/nafa.79.0001

126. North American Breeding Bird Survey. 2017. Available: https://www.pwrc.usgs.gov/bbs/.

127. eBird T. Global Big Day—8 May 2021 - eBird. 2021 [cited 23 May 2021]. Available: https://ebird.org/ebird/news/global-big-day-8-may-2021

128. Cornell Lab’s Citizen Science Projects. In: Citizen Science [Internet]. [cited 23 Aug 2021]. Available: https://www.birds.cornell.edu/citizenscience/about-the-projects/

129. Christmas Bird Count. In: Audubon [Internet]. [cited 23 May 2021]. Available: https://www.audubon.org/conservation/science/christmas-bird-count

130. About the Great Backyard Bird Count. In: Audubon [Internet]. 2015 [cited 23 May 2021]. Available: https://www.audubon.org/conservation/about-great-backyard-bird-count

131. GBIF - Global Biodiversity Information Facility. 2020 [cited 22 May 2020]. Available: https://www.gbif.org/

132. VertNet. [cited 23 Aug 2021]. Available: http://vertnet.org/

133. BISON - Biodiversity Information Serving Our Nation. [cited 23 Aug 2021]. Available: https://bison.usgs.gov/#home

134. C. Vega G, Pertierra LR, Olalla-Tárraga MÁ. MERRAclim, a high-resolution global dataset of remotely sensed bioclimatic variables for ecological modelling. Scientific Data. 2017;4. doi:10.1038/sdata.2017.78

135. von Storch H, Feser F, Geyer B, Klehmet K, Li D, Rockel B, et al. Regional reanalysis without local data: Exploiting the downscaling paradigm: Regional Reanalysis by Downscaling. J Geophys Res Atmos. 2017;122: 8631–8649. doi:10.1002/2016JD026332

136. Sen Gupta A, Tarboton DG. A tool for downscaling weather data from large-grid reanalysis products to finer spatial scales for distributed hydrological applications. Environmental Modelling & Software. 2016;84: 50–69. doi:10.1016/j.envsoft.2016.06.014

137. SEDAC - Socioeconomic Data and Applications Center. [cited 23 Aug 2021]. Available: https://sedac.ciesin.columbia.edu/theme/remote-sensing/data/sets/browse

138. Earthdata - NASA’s Earth Science Data Systems (ESDS) Program. [cited 23 Aug 2021]. Available: https://earthdata.nasa.gov//

139. NatureServe. [cited 23 Aug 2021]. Available: https://www.natureserve.org/

140. Araújo MB, Guisan A. Five (or so) challenges for species distribution modelling. Journal of Biogeography. 2006;33: 1677–1688. doi:10.1111/j.1365-2699.2006.01584.x

141. Peterson AT, Nakazawa Y. Environmental data sets matter in ecological niche modelling: an example with Solenopsis invicta and Solenopsis richteri. Global Ecology and Biogeography. 2008;0: 071113201427001-??? doi:10.1111/j.1466-8238.2007.00347.x

142. Smith AB, Santos MJ. Testing the ability of species distribution models to infer variable importance. Ecography. 2020;43: 1801–1813. doi:10.1111/ecog.05317

143. Barbet-Massin M, Jetz W. A 40-year, continent-wide, multispecies assessment of relevant climate predictors for species distribution modelling. Heikkinen R, editor. Diversity and Distributions. 2014;20: 1285–1295. doi:10.1111/ddi.12229

144. Warren DL, Matzke NJ, Iglesias TL. Evaluating presence-only species distribution models with discrimination accuracy is uninformative for many applications. Journal of Biogeography. 2020;47: 167–180. doi:10.1111/jbi.13705

145. Hamon J. Optimisation combinatoire pour la sélection de variables en régression en grande dimension : Application en génétique animale. Theses, Université des Sciences et Technologie de Lille - Lille I. 2013. Available: https://tel.archives-ouvertes.fr/tel-00920205

146. ABoVE - Arctic-Boreal Vulnerability Experiment. 2021 [cited 27 May 2021]. Available: https://above.nasa.gov/index.html?

147. Carroll ML, Loboda TV. The sign, magnitude and potential drivers of change in surface water extent in Canadian tundra. Environ Res Lett. 2018;13: 045009. doi:10.1088/1748-9326/aab794

148. Carroll ML, Schnase JL, Gill RL, Tamkin GS, Li J, Maxwell TP, et al. MERRA/Max: A Machine Learning Approach to Stochastic Convergence with a Multi-Variate Dataset. IGARSS 2020 Virtual Symposium. IEEE; 2020. Available: https://igarss2020.org/view_paper.php?PaperNum=4013

